# Decoding RNA Triple Helices: Identification from Sequence and Secondary Structure

**DOI:** 10.1101/2025.10.01.679706

**Authors:** Margherita A.G. Matarrese, Michela Quadrini, Nicole Luchetti, Federico Di Petta, Daniele Durante, Monica Ballarino, Letizia Chiodo, Luca Tesei

## Abstract

The discovery of long non-coding RNAs has revealed additional layers of gene-expression control. Specific interactions of lncRNAs with DNA, RNAs, and RNA-binding proteins enable regulation in both cytoplasmic and nuclear compartments; for example, a conserved triple-helix motif is essential for MALAT1 stability and oncogenic activity. Here we present a secondary-structure–based framework to annotate and detect RNA triple helices. First, we extend the dot–bracket formalism with a third annotation line that encodes Hoogsteen contacts. Second, we introduce TripleMatcher, which searches for a triple-helix pattern, filters candidates by C1^′^–C1^′^ distance thresholds, and merges overlaps into region-level zones. Using telomerase RNAs and RNA-stability elements with experimentally established triple helices (8 RNAs), TripleMatcher localized all annotated regions (structure-wise detection 8/8); geometric filtering removed most spurious candidates and improved precision (PPV from 0.42 to 0.81) and overall accuracy (F_1_ from 0.42 to 0.62) while maintaining sensitivity. Benchmarking eight predictors showed that pseudoknot-aware methods most reliably reproduce the local architecture required for detection, aligning secondary-structure quality with downstream triple-helix recovery. Applied prospectively, the framework identified candidate regions directly from predicted secondary structures and scaled to a screen of 4,147 RNAs, where distance filtering reduced 150,948 raw candidates to 90 geometrically feasible regions across seven molecules, including human telomerase complexes. Together, the notation and TripleMatcher provide a concise route from secondary structure to a small, interpretable set of triple-helix candidates suitable for targeted experimental validation.

## Introduction

RNA molecules adopt diverse secondary structures, such as stem-loops, bulges, G-quadruplexes, and pseudoknots, that regulate biosynthesis, stability, localization, and molecular interactions [1–3]. Experimental and computational approaches have linked structures to function [4–7]; chemical probing, NMR, and comparative analysis have clarified folding and dynamics [8]. Yet, capturing how these structures adapt to varying cellular conditions and how they modulate interactions with RNA or protein partners remains a significant challenge.

The role of secondary structures is particularly crucial for non-coding RNAs (ncRNAs), whose functionality extends beyond their linear nucleotide sequences and lack of protein-coding capacity. lncRNAs can act as scaffolds, bringing proteins and RNAs into proximity within nuclear and cytoplasmic compartments [9]. Deciphering the intricate folding patterns of these RNAs is essential for understanding their biological functions and could open avenues for therapeutic interventions in various diseases.

A significant example is the polyadenylated nuclear (PAN) RNA from *Kaposi’s sarcoma-associated herpesvirus* (KSHV); PAN RNA is abundant during the lytic phase, representing a large fraction of the cell’s polyadenylated RNA [10, 11]. Its expression and nuclear retention element (ENE) form a triple helix with the poly(A) tail via U•A–U interactions, protecting PAN from exonucleolytic decay [12–15] Another paradigmatic case is the Metastasis-Associated Lung Adenocarcinoma Transcript 1 (MALAT1), a multifunctional lncRNA with diverse roles in health and disease [16]. Conserved structural domains mediate interactions and nuclear localization [16, 17], and a conserved triple helix underlies its stability and nuclear accumulation [18, 19].

Beyond lncRNAs, triple helices are widespread in viral RNAs, riboswitches, catalytic ribozymes, and telomerase [17]. First observed by Felsenfeld et al. (1957) as U•A– U Hoogsteen triples and later confirmed in tRNA and telomerase RNA [20, 21], they contribute to RNA stability and regulation. In telomerase RNA, conserved U•A–U triples stabilize the pseudoknot required for catalytic activity [22– 25], and disrupting these triples impairs telomerase function [23, 26, 27]

Despite their biological significance, triple helices remain difficult to detect experimentally, primarily relying on crystallographic and NMR studies, with computational methods largely absent. Furthermore, standard notations for RNA secondary structure do not easily represent triple helices, complicating computational analyses [28, 29].

Here, we introduce a computational framework to characterize RNA triple helices. First, we extend the dot-bracket notation to encode Hoogsteen interactions. Then, we implement a four-stage workflow: (i) structural characterization from experimentally determined 3D RNA structures; (ii) secondary-structure prediction; (iii) development and validation of `TripleMatcher`, a search tool to detect putative triple-helix regions; (iv) identification and reliability assessment of triple helices.

Our ultimate goal is to enable the identification of tertiary structural motifs directly from nucleotide sequence via predicted secondary structure. Integrating these higher-order interactions may help bridge the gap between experimental and predicted RNA structural analyses, particularly for lncRNAs, where experimentally resolved structures are limited.

## Data and Methods

### Validation Dataset

We selected RNAs with experimentally determined structures containing triple helices, focusing on telomerase RNAs and two lncRNAs, MALAT1 and PAN (Fig. 1 and Table 1 of Supplementary Material). Major groove interactions were prioritized; minor groove interactions and intermolecular triple helices (e.g., NEAT1 [30]) were excluded.

**Table 1.** Matcher and 3DFilter command line options and their descriptions. For further details, see [54].

| Matcher Option | Description | Default Value |
| --- | --- | --- |
| -n <unpaired-nucleotide> | Nucleotide type for third strand (U, A, C, or G) | U |
| -b <canonical-base-pair> | Canonical base pair type (AU, UA, GC, or CG) | UA |
| -ml <pattern-minimum-length> | Minimum number of consecutive matches to detect | 4 |
| -st <sequence-tolerance> | Allowed mismatches/insertions/deletions in the unpaired sequence | 1 |
| -bt <base-pair-tolerance> | Allowed mismatches/insertions/deletions in the base-pair sequence | 1 |
| -pt <paired-tolerance> | Maximum number of base-paired nucleotides in unpaired region | 1 |
| -ct <consecutive-tolerance> | Maximum allowed interruptions in base-pair continuity | 1 |
| -p <pseudoknot-tolerance> | Maximum allowed bonds that are not part of a pseudoknot | Optional |
| -a <allow-all-submatches> | Find all sub-matches of exact bond matches | Optional |
| 3DFilter Option | Description | Default Value |
| -t <tolerance> | Tolerance in Ångström to be added to a maximum default distance 11 | 0 |

**Fig. 1.**
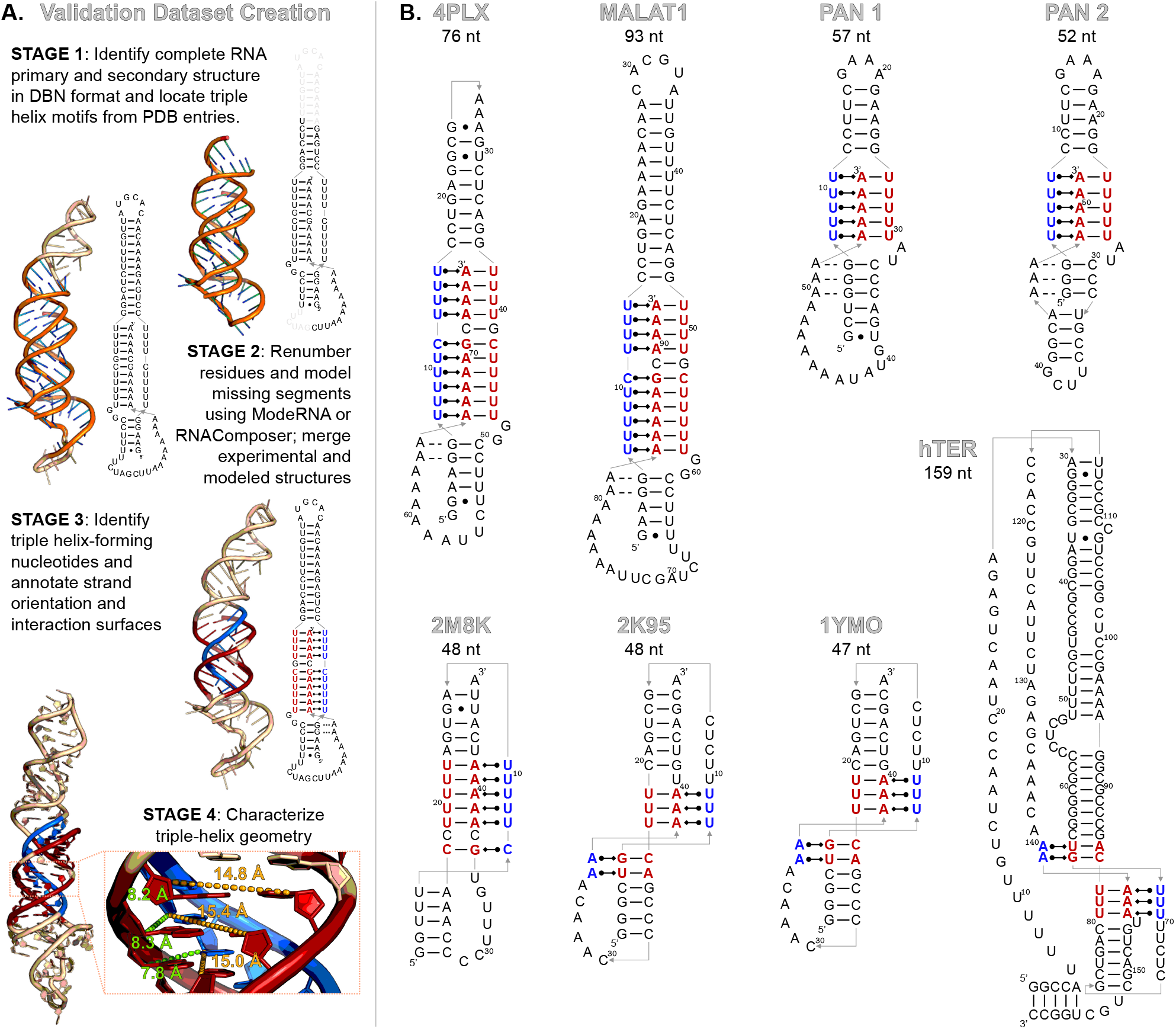
A) Three-stage workflow for preparing and annotating RNA triple helices. primary and secondary structures were extracted from reference publications and manually verified by two independent curators; only RNAs with an uninterrupted nucleotide sequence and an experimentally resolved triple-helix region in the PDB were retained. Residues were renumbered sequentially, missing segments were modeled with ModeRNA or RNAComposer and merged with experimental coordinates to obtain complete hybrid models. Nucleotides forming the triple helix were annotated from the secondary structure and 3D model, identifying the major-groove face of the WCF base-paired region and the orientation of the third strand. Example shown for MALAT1, where the resolved core was complemented by modeled flanking segments. After creating the final hybrid models, the triple helix is geometrically characterized by measuring C1^*′*^–C1^*′*^ distances from each third-strand nucleotide to the WCF base-paired region on (*i*) the interacting major-groove face (*First*) and (*ii*) the opposite face (*Second*). Right: overall 3D structure (third strand, blue; WCF region, red). Left: zoom of the triple-helix region with green dashed lines illustrating First distances (shorter) and yellow dashed lines illustrating Second distances (longer). **B) Secondary structures for the validation set** with triple-helix nucleotides highlighted: red, WCF base-paired region (major-groove face); blue, third strand engaged in Hoogsteen interactions.

MALAT1 is a nuclear lncRNA involved in gene regulation and cancer progression [31, 32]. Its 3’ end forms a bipartite triple helix composed of two runs of U•A–U base triples, interrupted by a C•G–C triplet and a C–G doublet that induce a ‘helical reset”, realigning the strands and preventing steric clashes [19]. Two A-minor interactions engage adjacent G–C base pairs [31, 32]. The core region is in the `4PLX` PDB [31, 33]; the full-length 3’ structure (upstream stem-loop and mascRNA tail) has been modeled to reconstruct missing residues using ModeRNA [34].

PAN RNA from KSHV forms two unimolecular triple helices [15]. The “PAN core triple helix” features a shortened apical P2 helix, designed to promote triple helix formation and stability, with mild protection against exonucleolytic degradation [14, 15]. The engineered “GCPAN triple helix” adds a GC clamp at the ENE base, anchoring the A-rich sequence to the lower base-paired region and increasing exonuclease resistance [15]. High-resolution PDBs for these unimolecular conformations are not available; the 3D models here derive from `3P22` and `6X5N`, integrating missing residues via ModeRNA [34].

Telomerase RNA (TER) provides the scaffold for ribonucleoprotein assembly and the template for telomere elongation. TER is conserved and contains essential features, including a conserved triple helix in the pseudoknot that supports catalytic activity and stability [35–38]. In *K. lactis*, the pseudoknot junction forms a triple helix stabilized by C•G– C and U•A–U base triples, with bound divalent cations [36]. In *H. sapiens*, the wild-type pseudoknot (`2K95`) includes a U41 bulge (U177 in [37]) that modulates catalysis; the deletion mutant (`2K96`/`1YMO`) lacks U177 but shares the network of base triples and a minor-groove A•G–C triple. For this study, the entire hTER core domain sequence (Fig. 2A from Zhang et al. [21]) was modeled by combining experimental elements and computational approaches. A full-length model of hTER was first generated using RNAComposer [39], which provided the global fold. The high-resolution triple-helix region was then inserted by replacing the corresponding segment with the experimentally resolved `2K95` structure, after superimposing and aligning the C1^*′*^ atoms of the nucleotides in common. Residues at the junction, connecting modeled and experimental regions, were refined using ModeRNA [34] to ensure smooth continuity and correct geometry (Fig. 1A).

**Fig. 2.**
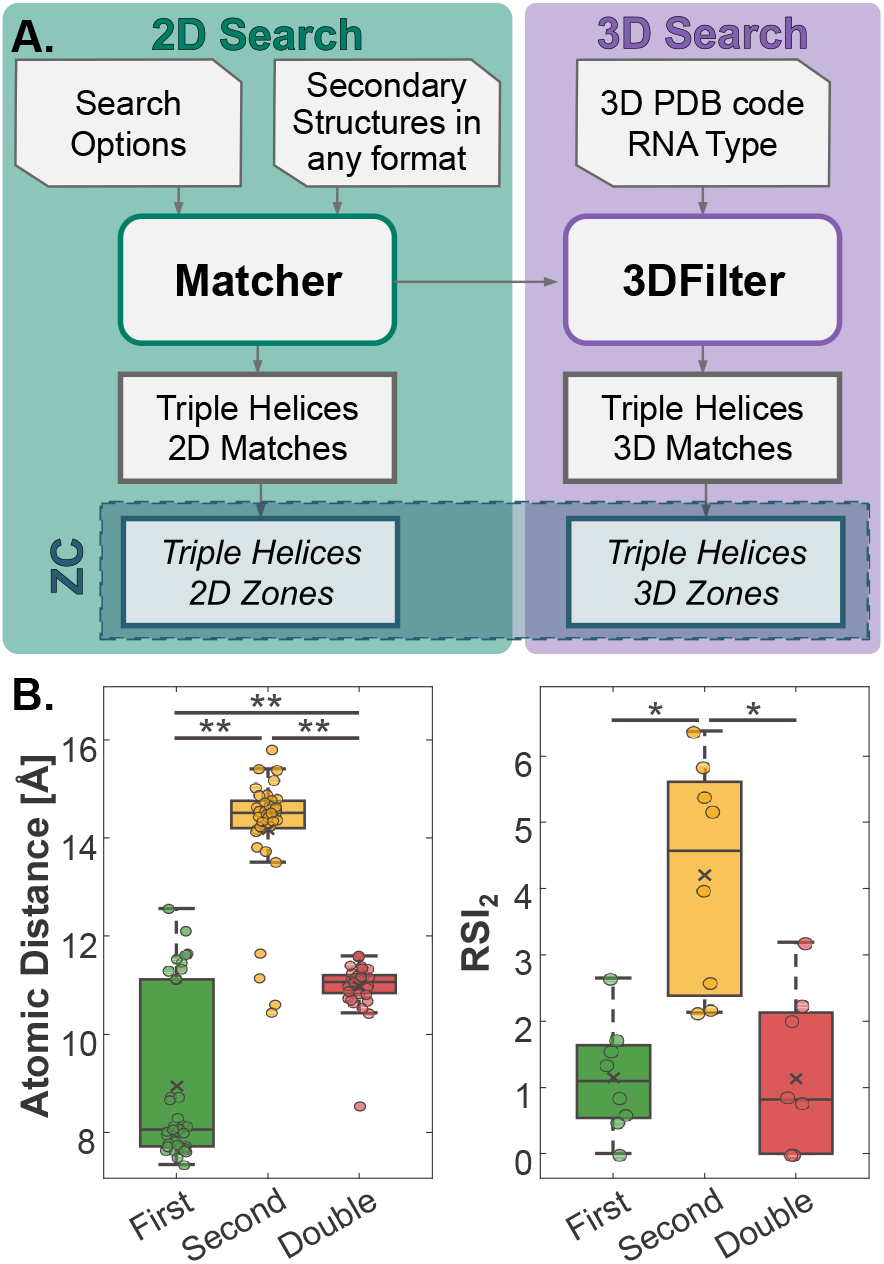
A) Schematic representation of the TripleMatcher tool. The Matcher scans RNA primary and secondary structures to report 2D-matches; the 3DFilter applies C1^*′*^–C1^*′*^ distance thresholds to keep only geometrically feasible triples (3D-matches); the ZoneCombiner (ZC) merges overlapping matches into non-overlapping zones. **B)** C1^*′*^–C1^*′*^ distance and RSI_2_ distributions for *First* (third-strand to the interacting WCF base on the major-groove face), *Second* (third-strand to the opposite WCF base), and *Double* (canonical WCF pairs). The separation between categories motivated the 3DFilter cutoff (default 11 Å) used to discard geometrically infeasible candidates. Each point denotes one base pair.

### Triple Helices Characterization

#### Secondary Structure Validation

Experimental secondary structure was obtained from the corresponding reference publication. When available, the PDB structure was parsed to extract the nucleotide sequence and secondary structure using RNAView through the RNAPDBee tool [40] (Fig. 1A). These structures were manually refined to match the reference models reported in the literature. We annotated all base pairs and third-strand nucleotides involved in triple-helix formation. When applicable, we marked the specific major-groove face where the third strand engages in Hoogsteen base pairing (Fig. 1B).

#### Augmented Dot-bracket Notation for Hoogsteen Pairs

With dot-bracket notation [29], pseudoknots can be represented using different types of brackets, but non-canonical interactions, such as Hoogsteen base pairs, are not encoded. To address this limitation, we introduce an augmented dot-bracket notation by adding a third string where (*i*) unpaired nucleotides involved in Hoogsteen pairing are marked with lowercase letters (e.g., `z, x, y, w, v`), and (*ii*) the facing nucleotides in the Watson– Crick–Franklin (WCF) base-paired region are marked with the matching uppercase letters (`Z, X, Y, W, V`). If a third-strand nucleotide interacts with both bases of a WCF pair, the uppercase letter is placed at both positions. All nucleotides not involved in Hoogsteen interactions are marked with a dash (`-`). All augmented notations for the Validation Dataset are reported in Table 4 of the Supplementary Material.

#### Three-dimensional Atomic Distance Assessment

For each base triple, we measured the Euclidean distance (in Å) between the C1^*′*^ atom of the third-strand nucleotide and the C1^*′*^ atoms of each nucleotide in the corresponding WCF base pair. To quantify geometric consistency within RNA triple helices, we computed a localized version of the Reconstruction Similarity Index (RSI_2_) [41], based only on the distribution of C1^*′*^–C1^*′*^ distances between base pairs forming the triple helix.

Lower RSI_2_ values indicate high geometric regularity of the triple helix, while higher values reflect structural divergence relative to typical base-pair spacing.

### Secondary Structure Prediction

For each RNA in the validation dataset, we predicted secondary structures using eight different folding tools: `CentroidFold` [42], `IPknot++` [43], `Mfold` [44], `pKiss` [45], `RNAfold` [46], `RNAshapes` [45], `RNAstructure` [47], and `vsfold5` [48].

Predicted structures were compared against experimentally refined secondary structures using standard evaluation
metrics: True Positive Rate (TPR), True Negative Rate (TNR), Positive Predictive Value (PPV), Fowlkes–Mallows index (FM), Matthews Correlation Coefficient (MCC), and Accuracy (ACC) [41]. In addition, for each predicted structure *p* and its corresponding experimental structure *e*, we computed the ASPRA [49, 50] and SERNA distances [51, 52], denoted respectively by *D*ASPRA (*p, e*), and *D*SERNA (*p, e*). Both distances quantify the dissimilarity between abstractions of secondary structures by considering only base pairs and measuring the minimal number of structural operations (i.e. insertions, deletions, and substitutions of base pairs) required to align two structures. *D*ASPRA (*p, e*) is based on tree alignment of structural trees, while *D*SERNA (*p, e*) adapts edit distance to structural sequences. We normalized both distances to the range [0, 1], using the maximum value observed across predictions for each tool 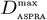 and 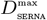. From these, we derived *normalized similarities* for each structure *p*, defined as:

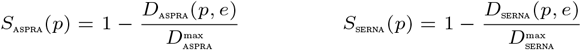

Metrics were computed per RNA and averaged within telomerase and stability element RNAs. This grouping was used to evaluate whether certain folding tools perform better depending on RNA class. Predicted structures, corresponding 2D matches, and the data used for metric computation are available on `Zenodo` [53].

### Triple Helices Identification

#### TripleMatcher Architecture

We developed `TripleMatcher`, a Java-based tool composed of two modules: the `Matcher` and the optional `3DFilter` [54]. The `Matcher` scans secondary structures to detect regions consistent with triple-helix motifs, using patterns derived from experimentally validated examples. These motifs typically involve a sequence of canonical WCF base pairs (e.g., A– U or C–G) and a corresponding run of unpaired nucleotides (commonly U or C) capable of forming Hoogsteen interactions. Fig. 2A shows the architecture of the tool.

The `Matcher` uses a dynamic programming algorithm to perform approximate pattern matching, producing a list of all identified 2D-matches. A 2D-match is defined as a pair of regions: one containing unpaired nucleotides that could serve as a third strand, and the other containing consecutive WCF base pairs (see Table 2 of Supplementary Material). Consecutive pairs must occur between adjacent nucleotide positions. Both regions must satisfy constraints on minimum length, pairing continuity, and the number of tolerated mismatches, insertions, or deletions, as specified by the user-defined options (see Table 1). By default, when an exact match is found, sub-matches are not reported to limit the number of 2D-matches generated. This behavior can be modified for bond matches by using the `-a` option.

When 3D structures are available, the `3DFilter` checks spatial feasibility by computing the atomic distances between the third strand and the matched WCF base pairs (see Table 3 of Supplementary Material), discarding matches that exceed a defined distance threshold. This ensures that only geometrically plausible triple-helix candidates, i.e., 3D-matches, are retained for downstream analysis.

Both 2D and 3D-matches can be aggregated into nonoverlapping zones by the independent `ZoneCombiner (ZC)` module. ZC merges sequences of matched unpaired nucleotides and WCF base pairs when they overlap within the same RNA molecule, reducing redundancy and highlighting broader structural regions likely to host a triple helix, even when detected as several nearby matches by the `Matcher` or `3DFilter`.

#### TripleMatcher Validation

We evaluated `TripleMatcher` by comparing its predicted 2D-matches against the experimentally annotated base triples for the validation dataset. Let *T* denote the set of annotated base triples (consisting of a WCF base pair and one nucleotide from a third strand forming a Hoogsteen interaction) in a given RNA. For each base triple *t ∈ T*, we define a true positive (TP) if there exists at least one predicted 2D-match in which all three nucleotide positions involved in *t* are present. If no such match exists for *t*, we record a false negative (FN). Conversely, any predicted 2D-match that does not fully cover any base triple in *T* is counted as a false positive (FP). We do not define true negatives (TNs), since we cannot exhaustively list every RNA segment incapable of forming a triple helix.

We quantified the *localization accuracy* (LA) by comparing the centres of the predicted and annotated regions. Let the RNA length be *N* . For the annotated triple helix *r* and a predicted 2D-match *m*, let *U*_*r*_, *U*_*m*_ be the sets of nucleotide indices belonging to the third strand and *D*_*r*_, *D*_*m*_ the sets of indices belonging to the WCF double strand. For *x ∈ {r, m}* we define the normalized centres:

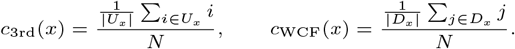

The LA score of *m* relative to *r* is then

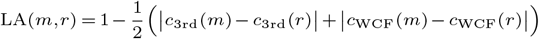

so that LA(*m, r*) *∈* [0, 1]; LA = 1 corresponds to perfect alignment of predicted and annotated centres, and smaller values indicate poorer localization.

Since the `Matcher` does not test spatial feasibility, we re-scored its output after applying the optional `3DFilter`. After filtering, a 3D-match can contain fewer base triples than the original 2D-match or be excluded entirely if no geometrically valid triples remain. To evaluate the impact of the `3DFilter`, we applied the same base-triple-level procedure used for the `Matcher`, defining: (*i*) TP as a retained predicted base triple overlapping an annotated one; (*ii*) FN as an annotated base triple with no retained match; and (*iii*) FP as a retained predicted triple with no corresponding annotation.

To quantify performance for both `Matcher` and `3DFilter`, we computed the following metrics: (*i*) PPV, measuring the proportion of predicted base triples that are correctly identified; (*ii*) TPR, measuring the fraction of true base triples recovered; (*iii*) FM index, the geometric mean of PPV and TPR; (*iv*) F1-score, the harmonic mean of PPV and TPR (*v*) localization accuracy, defined as the average offset between the center of a predicted match and the center of the annotated region; and (*vi*) `3DFilter` efficiency, defined as the percentage of FP base triples removed by the filter while retaining all TP.

#### TripleMatcher Usage – Search Dataset

We applied `TripleMatcher` to a large-scale search dataset composed of RNA secondary structures obtained from the `PhyloRNA database` at https://bdslab.unicam.it/phylorna[55] and from the `PDBe` archive [56]; it is available on `Zenodo` [53]. We used a total of 4,147 RNA structures, of which 2,207 contain pseudoknots. The dataset includes ribosomal RNAs (2897), transfer RNAs (1208), telomerase RNAs (19), and group I and II introns (23). We selected structures with available three-dimensional models to enable full usage of the `3DFilter` module.

The `Matcher` module scanned each RNA secondary structure for patterns consistent with triple-helix motifs, based on observed combinations of WCF base pairs and unpaired nucleotide sequences. Then, the `3DFilter` module was used to verify the spatial feasibility of each match.

#### TripleMatcher Usage – Predicted Secondary Structure

To assess the applicability of `TripleMatcher` in a fully computational setting, we applied the tool to predicted structures (Section Secondary Structure Prediction) per RNA in the validation dataset.

Predicted 2D-matches were evaluated against the annotated triple-helix regions using the same base-triple-level criteria described above. We computed the following metrics: (*i*) the number of TPs recovered per folding algorithm; (*ii*) the false positive rate (FPR), defined as the number of predicted base triples outside the annotated region; and (*iii*) structure-wise detection rate, defined as the fraction of RNAs for which at least one TP was recovered.

### Statistical Analysis

We used the Wilcoxon signed-rank test to compare atomic distances and RSI_2_ values across three categories of interactions in RNA triple helices: (*i*) between the third strand and the interacting side of the WCF double strand (*First*), (*ii*) between the third strand and the opposite side of the major groove (*Second*), and (*iii*) between nucleotides within the canonical WCF base pair (*Double*). Each base triple provided one measurement per category.

For evaluation metrics (TP, FP, TPR, PPV, F1-score, FM index, LA, and 3DFilter efficiency), per-RNA values were reported as median and interquartile range (IQR: 25th–75th percentile). Differences across groups, such as RNA type or folding tool, were assessed using the Wilcoxon rank-sum test. All statistical analyses were performed in MATLAB R2021b. Statistical significance was considered at *p <* 0.05 (*) and *p <* 0.001 (**).

## Results

### Triple Helices Characterization

We characterized the geometry of RNA triple helices using structural annotations and spatial distance metrics. Our augmented dot-bracket notation was used to annotate Hoogsteen interactions between a third strand and a canonical WCF double strand, allowing clear identification of triple-helix motifs within RNA secondary structures.

Fig. 2B shows the geometric analysis of base triples in the validation dataset. We measured C1^*′*^–C1^*′*^ distances across three categories: the WCF base engaged in Hoogsteen pairing (*First*), the opposite WCF base (*Second*), and standard WCF base pairs outside triple helices (*Double*). Distances on the Hoogsteen-interacting side were significantly shorter [8.5 (8.0– 11.2) Å] compared to the non-interacting side [15.0 (14.7–15.3) Å] and the WCF controls [10.4 (10.2–10.8) Å] (*p <* 0.001).

The corresponding RSI_2_ values supported this observation. RSI_2_ on the Hoogsteen-interacting face were low and similar to canonical WCF base pairs [1.7 (1.1–2.4) vs. 0.4 (0.3–2.1); *p* = 0.38], indicating structural regularity. In contrast, higher RSI_2_ values on the non-interacting side indicated greater spatial variability [4.8 (3.8–5.9); *p* = 0.008].

### Secondary Structure Prediction

Fig. 3A reports per-RNA metrics and the class aggregated values (telomerase, *n*=4; RNA-stability elements, *n*=4). Performance varied across both RNA types and prediction tools. Pseudoknot-aware methods (`pKiss`, `IPknot`, `vsfold`) outperformed the others on telomerase RNAs [MCC_pKiss_: 0.84 (0.52–0.95)], and among the methods without pseudoknot support the best was `mfold` [MCC_mfold_: 0.43 (0.28–0.57)]. Overall, `pKiss` was consistently top performing: on telomerase it showed (*i*) TPR 0.87 (0.74–0.97), (*ii*) TNR 0.94 (0.65–1.00), (*iii*) FM 0.92 (0.81–0.96), and (*iv*) ACC 1.00 (0.81–1.00); on RNA stability elements, metrics were: (*i*) TPR 0.93 (0.85– 0.95), (*ii*) TNR 0.83 (0.68–0.94), (*iii*) FM 0.93 (0.81–0.96), and (*iv*) ACC 1.00 (0.87–1.00). Non-pseudoknot algorithms often broke the pseudoknot stem or shifted its register, reducing MCC [median across `CentroidFold`, `mfold`, `RNAfold`, `RNAshapes`, `RNAstructure`: 0.34 (0.30–0.38)] while maintaining moderate TPR through over-pairing [0.66 (0.53–0.73)].

**Fig. 3.**
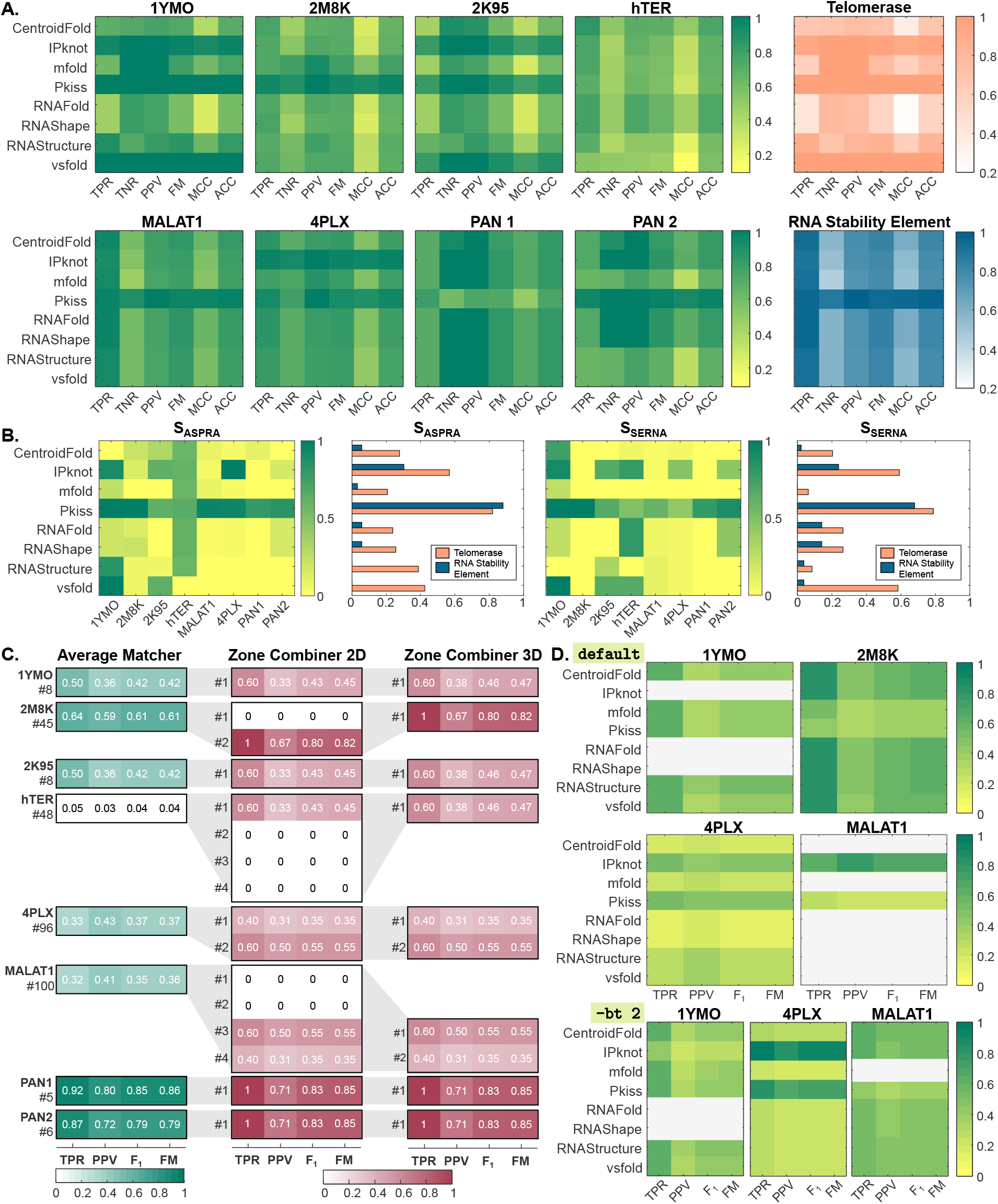
**A-B)** Performance of RNA secondary structure prediction algorithms on the validation dataset. Each matrix represents one RNA; rows are folding tools (CentroidFold, IPknot, mfold, pKiss, RNAfold, RNAshapes, RNAstructure, vsfold) and columns are evaluation metrics (TPR, TNR, PPV, FM, MCC, ACC). The two rightmost matrices show class averages (telomerase, top; RNA stability elements, bottom). Color encodes the score (0–1; light = low, dark = high). **C)** TripleMatcher performance by RNA and region. Rows list RNAs; columns report TPR, PPV, F_1_, and FM. For each RNA, the left panel (“Average Matcher”) shows averages across all raw 2D-matches; the number at left (e.g., #8 for 1YMO) is the count of raw 2D-matches used. The middle panel (“Zone Combiner 2D”) aggregates raw matches into non overlapping candidate regions (zones #1, #2, etc) and scores each zone over its base triple calls. Multiple zones indicate distinct candidate triple helix loci; in MALAT1, four zones arise because two alternative single strand segments and two separated base paired segments combine into four candidate triple helices. The right panel (“Zone Combiner 3D”) applies C1^*′*^–C1^*′*^ distance thresholds to remove geometrically infeasible triples; zones with all zeros contain no retained triples after filtering. **D)** Zone Combiner 2D on predicted secondary structures (four RNAs: (1YMO, 2M8K, 4PLX, MALAT1). Rows are folding tools; columns are base triple metrics (TPR, PPV, F_1_, FM). Gray cells indicate no detected region; metrics are undefined.

Abstract structure similarities mirrored these results (Fig. 3B). `pKiss` yielded the highest *S*_ASPRA_ and *S*_SERNA_, followed by `IPknot`, then `vsfold` (Fig. 3B). Class aggregates were higher for telomerase [*S*_ASPRA_: 0.33 (0.25–0.50), *S*_SERNA_: 0.26 (0.14–0.59)] than for RNA-stability elements [*S*_ASPRA_: 0.06 (0.02–0.18), *S*_SERNA_: 0.08 (0.03–0.19); *p*_ASPRA_ = 0.02, *p*_SERNA_ = 0.008]. Within telomerase, `hTER` (159 nt) showed the highest similarities [*S*_ASPRA_: 0.59 (0.58–0.61), *S*_SERNA_: 0.58 (0.03–0.79)], whereas the shorter constructs (`1YMO`, `2M8K`, `2K95`; 47–48 nt) were lower [*S*_ASPRA_ 0.25 (0.13–0.56), *S*_SERNA_ 0.18 (0.08–0.54)]. A plausible explanation is a length/context effect in the abstract-structure metrics. *S*_ASPRA_ and *S*_SERNA_ measure similarity from edits on *base pairs* only. `hTER` contains many helices outside the pseudoknot core that most tools predict correctly. These correct pairs dilute the impact of local errors in the pseudoknot, giving smaller edit distances and thus higher similarities. The shorter telomerase constructs concentrate most pairs within the pseudoknot. Any register shift or topology error then affects a large fraction of pairs and lowers similarity.

### TripleMatcher Validation

On the eight validation RNAs, the `Matcher` identified a median number of 26.5 (7–72) 2D-matches per RNA, for a total of 316 2D-matches. Per-RNA median counts were TP = 4 (3–4), FP = 4.5 (2–5), and FN = 2 (1.5–5.5); the structure-wise detection rate was 8/8 (100%). Per-RNA performance was TPR = 0.5 (0.32–0.75), PPV = 0.42 (0.36–0.66), F_1_ = 0.42 (0.36–0.70), FM = 0.42 (0.36–0.70), and LA = 0.97 (0.83–0.99). Class-wise summaries were comparable for telomerase vs. RNA-stability elements [TPR: 0.50 (0.27–0.57) vs. 0.60 (0.32–0.89), *p* = 0.63; PPV: 0.36 (0.20–0.47) vs. 0.57 (0.42–0.76), *p* = 0.08; F_1_: 0.42 (0.23–0.51) vs. 0.58 (0.36–0.82), *p* = 0.63; FM: 0.42 (0.23–0.52) vs. 0.58 (0.36–0.82), *p* = 0.63].

After applying the `3DFilter`, with default parameters, the number of retained 3D-matches decreased to 134, with a percentage reduction of 57.6%, reducing FPs while preserving TPR. PPV increased from 0.42 (0.36–0.66) to 0.81 (0.64–0.90) (Δ = +0.32, paired Wilcoxon *p* = 0.016), F_1_ from 0.42 (0.36– 0.70) to 0.62 (0.57–0.78) (Δ = +0.18, *p* = 0.023), and FM from 0.42 (0.36–0.70) to 0.65 (0.58–0.78) (Δ = +0.19, *p* = 0.023). LA improved from 0.97 (0.83–0.99) to 0.99 (0.91–0.99) (Δ = +0.04, *p* = 0.016). The filter removed 70% (68-82%) of FP base triples and retained all TPs in all the investigated RNAs.

Zone-level results are summarized in Fig. 3C. In the 2D aggregation (Fig. 3C central panel), most RNAs formed a single zone; `4PLX` had two, and `MALAT1` four, reflecting two alternative single-strand segments and two separated helical segments. After 3D filtering (Fig. 3C right panel), spurious zones were removed while the true triple-helix zones were retained. Specifically, in `2M8K`, two 2D zones collapsed to a single 3D-feasible zone with no improvement in terms of performances. `hTER` retained one zone with slightly improvement in performances, while the alternative 2D region was discarded. `4PLX` preserved two zones, and `MALAT1` reduced from four to two zones with the same respective scores, consistent with its bipartite triple helix. Consequently, `TripleMatcher` correctly localized all triple-helix regions in the validation set.

### TripleMatcher Usage: Screening and Identification of Putative Triple Helices

The `Matcher` applied to the search dataset filters 792 different PDBs, resulting in a total of 150,948 2D-matches. The `3DFilter` filters 62 different PDBs (full list on `Zenodo` [53]), for a total of 90 3D-matches. It identifies 7 distinct molecules, of which 1 is human telomerase RNA, 2 16S (from *E. coli* and *B. subtilis*), and 4 23S (from *E. coli, S. aureus, E. faecalis*, and *S. cerevisiae*). The human telomerase (PDBs `7V99` [57], `7QXA` and `7QXB` [58] from `PDBe`) contains the catalytic core, where the TERT protein interacts with the pseudoknot domain; indeed, a highly positively charged region on the surface of the TERT telomere repeat binding domain (TRBD) mediates intensive interactions with the major groove of pseudoknot stem P3 [57, 58], that contains the U•A–U triplets. The telomerase is indeed confirmed to contain the triple helix. In particular, despite the three structures being almost identical to the `2K95`, `1YMO`, and the `hTER` model included in our validation dataset, they were not identified by our first screening because the triple helix is classified as a pseudoknot in the reference papers. Interestingly, these telomerases are identified from PDB of protein-RNA complexes.

In the ribosomal RNA 16S of *B. subtilis* (`7QV1`), the region flagged by the `TripleMatcher` is not a triple helix; it corresponds to a junction. We see the same in 16S rRNA from *E. coli*: a region reported in 46 PDB entries shows three-base interactions that are best interpreted as a junction rather than a major-groove triple helix. In 23S rRNA, we also find non–triple-helix cases across 13 PDBs. In five structures (*E. coli* and *S. aureus*), a hairpin loop folds back onto its own helix (e.g., `6S12`). In six structures (*E. faecalis*), a loop from one hairpin contacts a separate helix. Finally, in two *S. cerevisiae* structures, a strand within a helix folds to create a local three-base interaction. These are junction-like or loop–helix contacts, not canonical triple helices.

### TripleMatcher Usage: Predicted Secondary Structure

We applied `TripleMatcher` to the secondary structures predicted by the eight tools (Section Secondary Structure Prediction), using the same defaults as in the validation. Across all predictions (8 RNAs *×* 8 tools = 64) predicted structures), the `Matcher` produced a total of 41 2D-matches; the fraction of predictions yielding at least one candidate triple helix region was 25/64 (39%). Per-prediction performance over predictions with at least one candidate was: (*i*) TPR = 0.50 (0.24–0.66), (*ii*) PPV = 0.33 (0.27–0.45), (*iii*) F_1_ = 0.43 (0.24–0.56), and (*iv*) FM = 0.45 (0.24–0.60). Pseudoknot-aware methods yielded candidates more often and with higher scores [detection rate 47%, F_1_ = 0.43 (0.29–0.54), FM = 0.45 (0.30–0.57)] than methods without pseudoknot support [31%, F_1_ = 0.39 (0.18–0.58), FM = 0.41 (0.18–0.62)].

By RNA, the proportion of tools producing at least one TP was highest for `2M8K` and `4PLX` (8/8), moderate for `1YMO` (5/8), and lower for `MALAT1` (only `IPknot` and `Pkiss`, Fig. 3D). Two edge cases are informative: for `PAN1`, `pKiss` returned a 2D-match with all predicted triples outside the annotated region; for `hTER`, `IPknot` likewise returned a 2D-match with TP=0, mapping away from the annotated site.

Model-specific defects in the predicted secondary structures explain several failures. In `2K95`, a single unpaired U41 breaks the adjacent paired stack in all tools; as a consequence, three annotated triples in that segment cannot be recovered, reducing the local TPs and prevent downstream detection at that site. In `PAN2`, enabling sub-matches (option `-a`) is required to recover a shorter, interrupted candidate with `Pkiss`; with this option, TP increases from 0 to 4 and *F*_1_ from 0 to 0.67. For `1YMO`, relaxing the base-pair tolerance to `-bt=2` enabled `IPknot` to recover the triple-helix locus (TP=2); with default settings, no candidate region was detected. For `4PLX`, the same relaxation compensated a register shift for both `pKiss` and `IPknot`, increasing TP from 5 to 8 (F_1_ from 0.48 to 0.73) and from 5 to 10 (F_1_ from 0.47 to 0.91), respectively. For `MALAT1`, setting `-bt=2` expanded the number of tools that reported a triple-helix candidate from 2 to 7, with a median *F*_1_ of 0.48 (Fig. 3D).

## Discussion

This work advances the computational analysis of RNA triple helices at two complementary levels. First, we extend the classical dot–bracket formalism with a third annotation line that captures Hoogsteen contacts, allowing triple-helical segments to be handled by any downstream parser that can read structured strings. Second, we implement `TripleMatcher`, a detector that (*i*) searches secondary structures for an empirically derived triple-helix pattern, (*ii*) filters the raw 2D-matches with atomic-distance thresholds (`3DFilter`) to rule out sterically impossible assemblies, and (*iii*) merges overlapping matches into higher-level “zones” (`ZoneCombiner`). In parallel, we quantitatively characterized triple-helix geometry on experimentally resolved references, using C1^*′*^–C1^*′*^ distance distributions and a localized RSI_2_ metric; these measurements both validate the motif and calibrate the `3DFilter` thresholds. Together, these elements provide a practical pipeline that starts from a primary sequence and ends with a short list of three-dimensionally plausible triple-helix candidate regions.

### Accuracy of the underlying secondary structures

Because `TripleMatcher` works on nucleotidic sequences and secondary structure, its success is bounded by the accuracy of that input. Our benchmarking confirms that pseudoknot-aware predictors, like `pKiss`, `IPknot`, and `vsfold`, best reconstruct the local architecture that triple helices require: a run of unpaired nucleotides positioned against a continuous WCF stack. Non-pseudoknot-aware methods, by contrast, frequently fragment the pseudoknot stem or shift its register, obliterating the geometric pre-conditions for Hoogsteen pairing (Fig. 3). Normalized pair-edit similarities (*S*_ASPRA_, *S*_SERNA_) make this point quantitatively and further reveal a length-dependent “dilution” effect: in long RNAs such as `hTER`, even substantial local errors have modest impact on global similarity, whereas in compact constructs every mis-paired base is proportionally more damaging. From a practical standpoint, this argues for combining experimental probing data or other constraints with folding algorithms whenever one aims to mine triple helices *in silico*.

### Performance of TripleMatcher on curated references

When supplied with experimentally refined secondary structures, `TripleMatcher` achieved a structure-wise detection rate of 100%, placing at least one true-positive match in every RNA of the validation set. The raw 2D search returned *∼*25 possible triple helices per molecule; geometric filtering discarded *∼*60% of them, halved the false-positive count, and raised PPV without loss of TPR. Zone aggregation (Fig. 3) distilled the remaining matches into one or two discrete loci per RNA, greatly easing manual inspection. In most cases, the surviving zone coincided with the annotated triple helix; where two zones persisted (`4PLX` and `MALAT1`), the duality reflected the genuine bipartite nature of the motif.

Beyond the curated set, several rRNA hits show why geometric screening and quick visual checks are necessary. In 16S rRNA from *B. subtilis* and *E. coli*, the flagged regions are junctions, and in 23S rRNA we see loop–helix or intra-helix contacts, all mimicking the 2D pattern but not canonical major-groove triple helices.

The `hTER` example illustrates an additional benefit; the second region flagged by the `Matcher` corresponds to the P2a helix [26], not part of the triple helix or a pseudoknot. This region accumulated more raw 2D-matches than the true triple helix before filtering (45 vs. 3), but was discarded by the `3DFilter`. Although it is not part of the triple helix, this apparent false positive points to a biologically important region of the `hTER` fold, as mutations in P2a/P2b reduce or abolish `hTER` activity [26].

### Fully computational use-case

Applying the same pipeline to purely predicted secondary structures proved feasible but more demanding. Only 39% of the 64 tool*×*RNA combinations produced any candidate region, and true positives were heavily skewed toward pseudoknot-aware predictions. `2M8K` and `1YMO` remained detectable in most predictors, whereas `PAN1` and `hTER` exemplified failure modes in which a 2D-match is found but lies outside the annotated triple helix.

Several misses could be rescued by modest parameter tweaks. Allowing sub-matches recovered a truncated but valid candidate in `PAN2`; relaxing the base-pair compensated for a two-base register shift in `1YMO`. Conversely, in `2K95` the apparent loss of three stacked pairs around U41 originates from the input sequence: this single nucleotide shifted the local pairing register and suppressed the three adjacent pairs used by the annotated triple helix, thereby precluding detection at that site. This underscores how even single-nucleotide changes in the supplied sequence can strongly bias secondary-structure prediction and, in turn, downstream motif detection.

## Conclusions and Future Directions

We developed an end-to-end pipeline that starts from an RNA sequence, uses its secondary structure (experimental or predicted), and outputs a small set of candidate triple-helix regions with quantitative descriptors. We validated it on two datasets with known 3D structures, and when 3D models are absent it still prioritizes likely triple-helix sites from predicted secondary structures.

Two factors primarily determine successful identification: (*i*) the quality of the secondary structure (notably, faithful recovery of pseudoknot topology and of a suitable run of unpaired nucleotides adjacent to a WCF stack), and (*ii*) sequence-intrinsic multiplicity of interaction-prone segments, which grows with RNA length and is therefore common in lncRNAs. The first can be mitigated by incorporating probing constraints or improved predictors; the second calls for stringent geometric screening and zone-level aggregation to control spurious patterns in long molecules.

`TripleMatcher` has the potential to suggest novel triple helix configurations that have not been previously characterized when applied to large-scale RNA structure datasets. As seen in the case of human telomerase, even if a triple helix is not present, it points out the presence of regions of interest for further investigations in terms of structural stability and RNA-RNA, DNA-RNA, or protein-RNA interactions.

Our rRNA screen underscores a practical caution: junctions and loop-mediated contacts can satisfy the 2D pattern; yet, they are not true triple helices. Pattern search should therefore be paired with geometric thresholds and a brief visual check, especially in long RNAs rich in junctions.

In future work, we will apply the pipeline at scale to curated ncRNA collections, with a focus on lncRNAs, and extend the analysis across species [59], prioritizing cases with established functional relevance. We will incorporate experimental constraints to improve secondary-structure fidelity and pursue targeted perturbations (e.g., CRISPR-based edits) to test whether predicted triple-helix modules influence RNA stability, localization, or macromolecular interactions *in vivo*. These studies will provide orthogonal validation of the *in silico* predictions and refine the rules that govern triple-helix formation in long RNAs.

## Key Points

1. We introduce TripleMatcher, a computational framework that detects RNA triple helices from sequence and secondary structure using rule-based structural matching on unpaired nucleotides and adjacent base-pair bonds.
2. TripleMatcher enhances detection precision through 3D spatial filtering, ensuring that predicted motifs correspond to physically plausible tertiary structures.
3. The framework enables large-scale screening of experimentally solved and computationally predicted RNAs, yielding a limited set of high-confidence candidate triple helices.
4. By bridging sequence, secondary structure, and tertiary geometry, TripleMatcher provides a generalizable tool for exploring higher-order RNA interactions and guiding experimental validation.

## Data Availability

The modeled RNA structures (`MALAT1`, `PAN1`, `PAN2`, `hTER`), curated secondary structures, and annotations are available on `Zenodo` [53]. Predicted structures, evaluation datasets, and metric computation files are also available on Zenodo. The `TripleMatcher` source code is open source at https://github.com/bdslab/triplematcher [54] with documentation and usage examples in the repository.

## Author Contributions

**MAGM**: Conceptualization, Data curation, Formal analysis, Investigation, Methodology, Software, Validation, Visualization, Writing - original draft, Writing - review & editing; **MQ**: Formal analysis, Data curation, Software, Writing - review & editing; **NL**: Formal analysis, Data curation, Visualization, Writing - review & editing; **FDP**: Formal analysis, Software, Validation, Writing - review & editing; **DD**: Writing - review & editing; **MB**: Conceptualization, Funding acquisition, Writing - review & editing; **LC**: Conceptualization, Funding acquisition, Data curation, Investigation, Supervision, Validation, Writing - original draft, Writing - review & editing; **LT**: Conceptualization, Funding acquisition, Software, Validation, Supervision, Project administration, Writing - original draft, Writing - review & editing.

## Biographical Notes

**Margherita A.G. Matarrese** is Assistant Professor of Biomedical Engineering at Campus Bio-Medico University of Rome. Her research includes biomedical signal processing, multimodal neuroimaging integration, mathematical and computational modeling of brain function, network theory, and RNA structure analysis.

**Luca Tesei** is Associate Professor of Computer Science at the University of Camerino, Italy. His research includes computational biology, bioinformatics methods, formal methods, and the analysis, abstraction, and comparison of RNA structures.

