## Supplementary Tables for "Decoding RNA Triple Helices: Identification from Sequence and Secondary Structure"

for the article *Decoding RNA Triple Helices: Identification from Sequence and Secondary Structure*

#### Overview

This supplementary material provides additional tables to support the analyses and methods described in the main article. It includes:

- Details about the RNAs in the Validation Dataset.
- Output of `TripleMatcher` for Triple Helix Identification: Examples of 2D and 3D Matches.
- Augmented dot-bracket notation for the Validation Dataset.

#### Validation Dataset Description

**Supplementary Table 1.** RNA Validation Dataset.

| RNA source | Base triples No. | Base triples Identity | Structure | PDB ID / Ref. |
|---|---|---|---|---|
| ***RNA stability elements*** | | | | |
| *Homo sapiens* MALAT1 ENE | 9 | U•A–U | X-ray, 3.10 Å | 4PLX [1] |
| | 1 | C•G–C | | |
| *Homo sapiens* MALAT1 | 9 | U•A–U | Modeled from 4PLX | [1] |
| | 1 | C•G–C | | |
| KSHV PAN RNA 1 | 5 | U•A–U | Modeled from 6X5N | [5] |
| KSHV PAN RNA 2 | 5 | U•A–U | Modeled from 3P22 | [4, 5] |
| ***Telomerase*** | | | | |
| *Kluyveromyces lactis* | 1 | C•G–C | NMR | 2M8K [2] |
| | 5 | U•A–U | | |
| *Homo sapiens* | 1 | A•U–A | NMR | 1YMO, 2K95 [6, 3] |
| | 1 | A•G–C | | |
| | 3 | U•A–U | | |
| *Homo sapiens* | 1 | A•U–A | Modeled from 2K95 | [7, 3] |
| | 1 | A•G–C | | |
| | 3 | U•A–U | | |


### Examples and Explanation of 2D– and 3D–matches of `TripleMatcher` tool

**Supplementary Table 2.** Example of two predicted 2D-matches from the `2M8K` RNA structure. The left column (`unpaired indices`) lists the positions of unpaired nucleotides predicted to form Hoogsteen interactions as the third strand. The right column (`WCF basepairs`) shows the corresponding WCF base pairs potentially involved in triple-helix formation. In the first row, the unpaired region lies outside the annotated triple helix; in the second row, it corresponds to the validated Hoogsteen strand. In both cases, the paired region belongs to a known triple helix.

| `indices_seq` | `indices_bond` |
|---|---|
| 31;32;33;34;35 | (17,42);(18,41);(19,40);(20,39);(21,38);(23,36) |
| 6;7;8;9;10;11 | (17,42);(18,41);(19,40);(20,39);(21,38);(23,36) |

**Supplementary Table 3.** Example of spatial distance data extracted from the RNA structure `2M8K`, corresponding to the second 2D-match listed in Supplementary Table 2. The values are taken from the `distance_info` column of the `TripleMatcher` CSV output and reorganized into three columns for improved readability. Each entry is a triple of the form ($index_1$; $index_2$; distance), where the distance (in Å) is the Euclidean distance between the C1′ atoms of the two nucleotides involved. **Interaction A** lists distances between the unpaired third-strand nucleotide and the base in the WCF pair that is *not* involved in Hoogsteen pairing. **Interaction B** shows distances between the third strand and the Hoogsteen-interacting base, which are expected to fall below the 11 Å threshold (average: 8.25 Å). **Interaction C** reports distances between the two nucleotides forming each canonical WCF base pair.

| **Interaction A** | **Interaction B** | **Interaction C** |
|---|---|---|
| (11; 17; 14.96) | (11; 42; 7.78) | (17; 42; 11.06) |
| (10; 18; 14.60) | (10; 41; 7.77) | (18; 41; 10.96) |
| (9; 19; 14.82) | (9; 40; 7.95) | (19; 40; 10.40) |
| (8; 20; 14.89) | (8; 39; 8.50) | (20; 39; 9.70) |
| (7; 21; 14.57) | (7; 38; 9.01) | (21; 38; 9.76) |
| (6; 23; 13.02) | (6; 36; 8.49) | (23; 36; 7.41) |

### Augmented Dot-bracket Notation for the Validation Dataset

**Supplementary Table 4.** Augmented dot-bracket notation for RNAs in the validation dataset. For each RNA, the sequence and its corresponding secondary structure are shown using standard and extended dot-bracket symbols (including pseudoknot brackets). Below each structure, a third annotation line encodes Hoogsteen interactions involved in triple helix formation. Each Hoogsteen pair is represented by a matching lowercase-uppercase letter pair (e.g., `zZ`), where the lowercase letter marks the unpaired nucleotide (third strand) and the uppercase letter marks its corresponding interaction site on the WCF pair.

| RNA | Augmented dot-bracket notation |
| --- | --- |
| 4PLX | GGAAGGUUUUUCUUUUCCUGAGGCGAAAGUCUCAGGUUUUGCUUUUUGGCCUUUCUUAAAAAAAAAAAAAGCAAAA<br>.(((((..........(((((((((..)))))))))[[[[[[[[[[[..)))))...........]]]]]]]]]]]<br>------zyxwvutsrq-------------------------------------------------ZYXWVU-TSRQ |
| MALAT1 | GAAGGUUUUUCUUUUCCUGAGAAAACAACACGUAUUGUUUUCUCAGGUUUUGCUUUUUGGCCUUUUUCUAGCUUAAAAAAAAAAAAAGCA<br>AAA<br>(((((..........((((((((((((((......)))))))))))))[[[[[[[[[[[..))))).................]]]]]]]]<br>]]]<br>-----zyxwvutsrq-------------------------------------------------------------------ZYXWVU-T<br>SRQ |
| PAN 1 | GCUGGGUUUUUCCUUCGAAAGAAGGUUUUUAUCCCAGUGUAUAAAAAAAAAAAAAAA<br>((((((.....(((((....)))))[[[[[..))))))..............]]]]]<br>------zyxwv-----------------------------------------ZYXWV |
| PAN 2 | GGGUUUUUUCCUUCGAAAGAAGGUUUUUAUCCCUGCCUUCGGGCAAAAAAAA<br>(((.....(((((....)))))[[[[[..)))((((....))))...]]]]]<br>---zyxwv---------------------------------------ZYXWV |
| 2M8K | GGUUUCUUUUUAGUGAUUUUUCCAAACCCCUUUGUGCAAAAAUCAUUA<br>(((((......[[[[[[[[[[.[))))).......].]]]]]]]]]].<br>-----zyxwvu------------------------Z-YXWVU------ |
| 2K95 | GGGCUGUUUUUCUCGCUGACUUUCAGCCCCAAACAAAAAAUGUCAGCA<br>((((((........[[[[[[[[[))))))........]]].]]]]]].<br>----ZYxwv--------------------------zyXWV-------- |
| 1YMO | GGGCUGUUUUUCUCGCUGACUUUCAGCCCCAAACAAAAAAGUCAGCA<br>((((((........[[[[[[[[[))))))........]]]]]]]]].<br>----ZYxwv--------------------------zyXWV------- |
| hTER | GGCCAUUUUUUGUCUAACCCUAACUGAGAAGGGCGUAGGCGCCGUGCUUUUGCUCCCCGCGGGCUGUUUUUCUCGCUGACUUUCAGCCCG<br>CGGAAAAGCCUCGGCCUGCCGCCUUCCACCGUUCAUUCUAGAGCAAACAAAAAAUGUCAGCAGCUGGCC<br>(((((........................(((((((((((.....((((((.....((((((((((........[[[[[[[[[)))))))<br>)))))))))....))))).)))))).........................]]].]]]]]]...)))))<br>----------------------------------------------------------------ZYxwv---------------------<br>-------------------------------------------------zyXWV--------------- |
